# GraphMHC: neoantigen prediction model applying the graph neural network to molecular structure

**DOI:** 10.1101/2023.08.25.554848

**Authors:** Hoyeon Jeong, Young-Rae Cho, Jungsoo Gim, Seung-Kuy Cha, Maengsup Kim, Dae Ryong Kang

## Abstract

Neoantigens are biomarkers that can predict the prognosis associated with immune checkpoint inhibition by estimating the binding potential of candidate peptides to somatic mutation and major histocompatibility complex (MHC) proteins. Although deep neural networks have been primarily used for these prediction models, it is difficult to consider the models reported thus far as accurately representing the interactions between biomolecules. In this study, we propose the GraphMHC model, which utilizes a graph neural network model through molecular structure to simulate the binding between MHC proteins and peptide sequences. Amino acid sequences sourced from the immune epitope database (IEDB) undergo conversion into molecular structures. Subsequently, atomic intrinsic informations and inter-atomic connections are extracted and structured as a graph representation. Bindings are classified by feeding them into the GraphMHC network, comprising stacked graph attention and convolution layers. The prediction results from the test set using the GraphMHC model showed a high performance with an area under the receiver operating characteristic curve of 92.2% (91.9-92.5%), surpassing the baseline model. Moreover, by applying the GraphMHC model to melanoma patient data from the Cancer Genome Atlas project, we found a borderline difference in overall survival and a significant difference in stromal score between the high and low neoantigen load groups. This distinction was not present in the baseline model. This study presents the first feature-intrinsic method based on biochemical molecular structure for modeling the binding between MHC protein sequences and neoantigen candidate peptide sequences. The model can provide highly accurate suitability information for cancer patients who want to apply immune checkpoint inhibitors.

**Author summary:** 

## Introduction

Cancer is the leading cause of death in a large number of countries worldwide [Sung et al., 2021] and the number one cause of death in South Korea. [Jung et al., 2020] Cancer mainly results from somatic mutations. [Anand et al., 2008] Cancer immunotherapy basically works by encouraging tumor cells to recognize the MHC protein as foreign, leading to activation of the cytotoxic T cell receptor (TCR) and CD8 coreceptor. However, the mechanism by which this occurs acts as an immune checkpoint, suppressing overactivation of the immune system, with proteins such as the programed cell death protein 1 (PD-1) and cytotoxic T lymphocyte antigen-4 (CTLA-4) the main actors in these pathways. Third-generation anticancer drugs block or inhibit this immune checkpoint. [Sharma and Allison, 2015] Despite these efforts, only a small number of subjects respond well to immune treatment, [Nam et al., 2019] and it is limited because of the high expense. [Ventola, 2017] Neoantigens, or neoplasmic antigens, are tumor-specific antigenic determinants or epitopes that consist of 9-mer-long peptides that are cleaved by proteasome internal organelles and act as biomarkers predicting immune checkpoint inhibition, [Jiang et al., 2019, Yi et al., 2018] which can be estimated by predicting the binding potential of major histocompatibility complex (MHC) proteins to candidate peptides associated with somatic mutations.

MHC proteins bind with peptides via non-covalent hydrogen bonds. The half maximal inhibitory concentration (IC_50_) between MHC and isolated peptides can be experimentally determined, [Sette et al., 1994] and the results of binding attempts can be found in the Immune Epitope Database (IEDB). [Vita et al., 2015] Prediction models comprising deep neural networks can utilize the information in the IEDB, and several studies have used NetMHCpan-4.0 [Jurtz et al., 2017] and 4.1 [Reynisson et al., 2020] to model bonding between MHC and peptide using these data. These models are representative de facto, but use conventional neural networks. MHCAttnNet uses a recurrent neural network (RNN) for string processing, [Venkatesh et al., 2020] DeepNeo uses a convolutional neural network (CNN), [Kim et al., 2020] and DeepImmuno uses CNN together with graph convolutional networks (GCN) [Kipf and Welling, 2016] to understand the relationships between amino acids rather than the constituent atoms. [Li et al., 2021] However, the fact that these models only attempt to connect vectors in one dimension or transfer them into two-dimensional matrices means that the biomolecular interactions cannot be accurately discerned. Meanwhile, studies have reported the use of the Simplified Molecular Input Line Entry System (SMILES) [Weininger, 1988] and graph neural networks (GNN), which is from an adjacent academic field and expresses molecular structure in the form of polymer interactions, as input for deep neural networks. This type of input corresponds to protein-protein interaction (PPI), drug-target affinity (DTA), and drug-drug interaction (DDI). Interaction methods can be grouped into two categories: concatenation [Yang et al., 2022, Nguyen et al., 2021, Öztürk et al., 2018] and coattention. [Nyamabo et al., 2021, Deac et al., 2019] However, since these graph-graph interaction can be combination between embedded vectors extracted from each graph rather than direct combination between the nodes constituting the two graphs, there is a limit to model the interaction between all components of the graph. This study proposes the GraphMHC model, which uses data from the IEDB and the GNN model to illustrate the binding between MHC proteins and peptide sequences via the molecular structure. The determination of the feature extracted from combining an MHC protein with a peptide as a neoantigen is performed through multi-layered graph convolution when using this method. The GraphMHC model was validated against the baseline model, [Reynisson et al., 2020] and applied to melanoma patient data from the Cancer Genome Atlas (TCGA) project. Data were divided into low and high neoantigen load groups for comparison of the clinical differences.

## Materials and methods

In this section, the methods used to build and validate the neoantigen prediction model GraphMHC are presented.

### GraphMHC: a neoantigen prediction model based on the GNN using MHC class I from the IEDB

The processes for building GraphMHC, which is a neoantigen prediction model based on GNNs, are presented in detail in this section. The overall pipeline can be seen in Figure 1.

**Fig 1.**
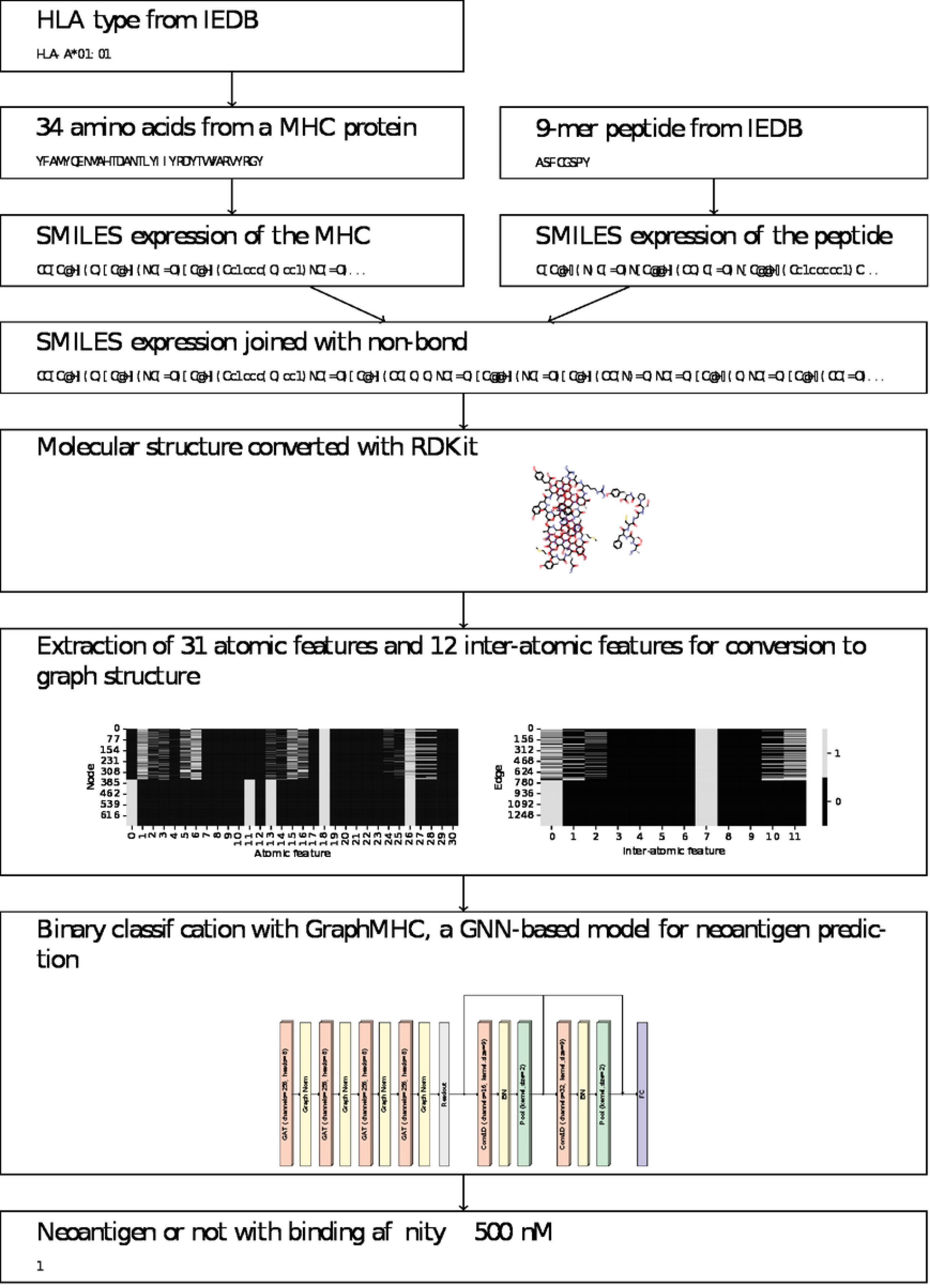
Overall pipeline for modeling neoantigen prediction. HLAs and 9-mer peptides from the IEDB are converted into molecular structures, and the GraphMHC model is used to classify probable neoantigens from the extracted atoms and interatomic features.

The Immune Epitope Database (IEDB), [Vita et al., 2015] a binding affinity data set for peptides and HLA types, was used to construct the model for predicting neoantigens. The MHC class I dataset was used for neoantigen prediction. The use of the MHC class II dataset is described separately in the section describing extra-validation.

Referring to previous studies, a result with a binding affinity of 500 nM or less was determined to be a neoantigen. [Łuksza et al., 2017, Vitiello and Zanetti, 2017, Richman et al., 2019]

#### Conversion from the molecular structure of amino acids to dataset of interatomic graph structures

In the IEDB dataset, 157,325 rows pertaining to humans were utilized. These rows were then transformed into amino acids that constitute the MHC protein, employing the conversion data from NetMHCpan-4.1. [Reynisson et al., 2020] Out of these, 157,084 were converted, excluding types for which conversion information was unavailable.

Classification was based on binding affinity, with binding assumed for IC_50_ ≤ 500 nM, and non-binding for IC_50_ *>* 500 nM. The dataset was split, with 80%, or 125 667, for training and 20%, or 31 417, for testing. The conversion process from database to dataset is as follows.

First, human leukocyte antigens (HLA) from the IEDB are converted to MHC amino acid sequences. Second, MHC and peptide sequences are converted using SMILES [Weininger, 1988] structures using the RDKit 2022.03.2 library. The expressions used are given in Table S1. Third, Join the two SMILES strings with non-bond notation together (.). Fourth, Convert to molecular structure using RDKit. Hydrogen atoms that were omitted from the symbol must be expressed at this point. The molecular structure is expressed in Figure S2. Fifth, Convert to graph structure using the RDKit library to encode vectors and matrices. The feature information of the graph representation is shown in Table 1. Each feature is encoded via one-hot encoding and constructed as a sparse matrix. The graph structure is expressed in Figure S2 using NetworkX 2.8.4 library with Kamada Kawai layout. [Kamada et al., 1989] Characteristics of graph representations describing bound and unbounded data are shown in the Table S2. Sixth, Converting graph dataset using the PyTorch Geometric (PyG) [Fey and Lenssen, 2019] 2.1.0 library.

**Table 1.** Components of graph representation.

| Graph representation | Components | Number of features |
| --- | --- | --- |
| Node feature vectors with one-hot encoding | Atom symbol (H, C, N, O, S), hybridisation (SP3, SP2, SP, S, SP3D, SP3D2), degree of covalent bonding, number of bonded hydrogen atoms, chirality (CCW, CW or others), aromaticity, inclusion in rings, formal charge, and number of radical electrons | The number of features per node is 31 |
| Edge indices (adjacency matrix) | Between the starting atom and the ending atom |  |
| Edge feature vectors with one-hot encoding | covalent bond type (single, aromatic, double, triple), stereo type (any, cis, E, none, trans, Z), ring bond, conjugate bond | The number of features per edge is 12 |

#### Architectural design of the GNN GraphMHC for graph classification

Layer-by-layer architecture of GraphMHC, the GNN model for MHC-peptide binding, is presented in a schematic diagram provided in Figure 2. In this model, graph attention [Bahdanau et al., 2014] was used as graph convolution, where the attention factor is multiplied to assign importance to nodes. Another noteworthy element of this model is the stacking of conventional one-dimensional convolution layers after graph convolutions. This procedure enables the re-extraction of a given feature vector multiple times for classification purposes. The stacking steps of the model are subdivided as follows.

**Fig 2.**
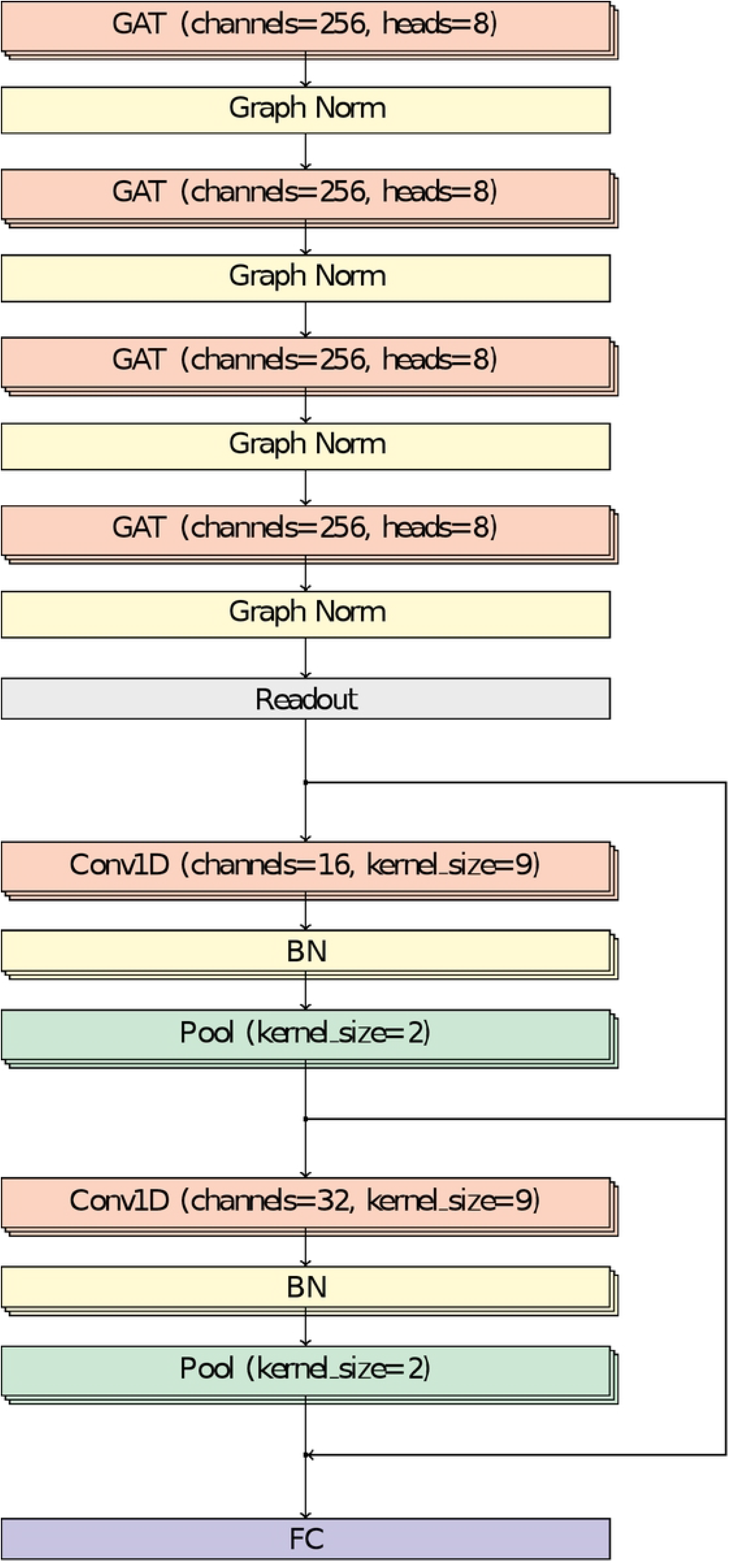
Architecture of the GraphMHC model, a GNN that predicts neoantigens via MHC-peptide binding information. Detour arrows between Conv1D layers mean that skip connection. Dropout was set at 0.1 for all layers. GAT: graph attention layer, FC: fully-connected layer, Graph Norm: graph normalization, Conv1D: 1-dimensional convolution, BN, batch normalization, Pool: average pooling.

First, stack four layers of graph attention [Veličković et al., 2017] using PyTorch Geometric (PyG) [Fey and Lenssen, 2019] 2.1.0 library. Second, for graph classification, the readout layer is the mean value of the node feature vectors. Third, two layers of 1-dimensional convolution are stacked with the PyTorch [Paszke et al., 2019] 1.11.0+cu113 framework. At this time, add a skip connection [He et al., 2016] between layers. Fourth, classify via sigmoid in fully-connected layer.

#### Model training and evaluation

ADAM [Kingma and Ba, 2014] was used as the optimizer for learning. For tensor calculation during the deep neural network model training, massive parallel processing was performed using RTX 3090 24GB graphic processing units (GPU).

Scikit-learn [Pedregosa et al., 2011] 1.1.2 was used to evaluate the classification model, and the 95% AUC-ROC confidence interval was obtained through the DeLong method [DeLong et al., 1988] in MedCalc [Schoonjans et al., 1995] 20.106. 95% confidence intervals for other metrics were calculated directly.

For comparison with models comprising fewer layers, two cases were constructed; one of which involved subtracting two of four GNN layers while the other involved the subtraction of two CNN layers.

### Neoantigen load predictions on next generation sequencing (NGS) data from The Cancer Genome Atlas database

The constructed model was applied to next-generation sequencing (NGS) data from actual cancer patients. Data were sourced from The Cancer Genome Atlas (TCGA). [Weinstein et al., 2013] Skin cutaneous melanoma (SKCM) data was used because of the effective response that this type of cancer has to immunotherapy. Data were primarily downloaded from the Broad Institute Firehose website.

Sequence information for normal samples, which is not disclosed by TCGA-SKCM, is required for HLA typing and was thus obtained from previous studies. [Coelho et al., 2020] The HLA types were reported to be Optitype [Szolek et al., 2014] and were converted to 34-mer amino acids using NetMHCpan-4.1. [Reynisson et al., 2020]

The procedure for translating the mutation into a 9-mer peptide and identifying it as a neoantigen is as follows.

First, convert from mutation annotation format (MAF) to variant calling format (VCF) using maf2vcf. [Park et al., 2021] Second, annotate as Variant Effect Predictor (VEP). [McLaren et al., 2016] At this time, the sequence information for the GRCh38 reference genome is used. Third, select only missense variants. Fourth, convert to amino acids with R customProDB. [Wang and Zhang, 2013] The chromosome nomenclature must be consistent for this process. Fifth, only the region in which the actual mutation appears is truncated to a length of 9-mer. [Kim, 2020] Sixth, convert the MHC protein sequence and peptide sequence together to form a graph structure. Seventh, predict the binding affinities using GraphMHC.

Thus, data from a total 310 subjects were used for predicting neoantigen load by combining 9-mer peptides that are considered neoantigen candidates with HLA type information. For more details of the selection process, see Figure S3. Candidates are then predicted neoantigens or not using GraphMHC.

#### Comparison of groups according to high and low neoantigen load

Two groups were formed around the median, with high and low neoantigen loads. Comparison was performed using survival analysis and immunity score analysis. Clinical information on survival was also obtained from the Firehose website. This study uses overall survival as the analysis target, for which Lifelines 0.27.0 was used.

Estimates of stromal cell score, immune cell score, and tumor purity were obtained from the ESTIMATE website.and utilized. [Yoshihara et al., 2013] Data were calculated from the TCGA expression data. The stromal score refers to the number of stroma cells within tumor tissue, [Hanahan and Weinberg, 2011, Kalluri and Zeisberg, 2006] and was used because stromal cells have been reported to be involved in tumor growth and disease progression. The immune score indicates the infiltration of immune cells into tumor tissues and is an immunological biomarker for prognosis prediction and therapeutic response. [Galon et al., 2012] The ESTIMATE score relates to tumor purity and is a combination of the stromal and the immune scores. Scipy 1.9.3 was used for the comparison test.

### Extra-validation of the model using MHC class II from the IEDB

Although neoantigens are related to MHC class I and CD8^+^, MHC class II and CD4^+^ have also been reported complementary, with less variation between patients. [Pyke et al., 2018, Sun et al., 2017] Several model studies have investigated this concept, [Reynisson et al., 2020, Shao et al., 2020] and a model with the same architecture as the proposed model was trained using MHC class II data from the IEDB for extra-validation in this study. The same classification threshold of 500 nM was used. Data with reduced similarity were selected, and training and test sets comprising 85 708 and 21 427 data points, respectively, were used after pre-processing.

### Baseline models for comparison

The model NetMHCpan-4.1 [Reynisson et al., 2020] was used for comparison. Classification using IEDB data and comparison between groups using TCGA-SKCM data were applied equally. NetMHCIIpan-4.0 was used for extra-validation. [Reynisson et al., 2020] In order to convert the obtained binding affinity to a value between 0 and 1, studies [Zhang et al., 2009, Andreatta and Nielsen, 2016, Jurtz et al., 2017] such as those involving NetMHCpan-4.0 used the expression 1 − log(·)*/* log(50000), whereas an equation 1*/*(1 + exp(·)) similar to the sigmoid was used for the same comparison in this study.

## Results

In this section, the GraphMHC model, which is proposed for neoantigen prediction based on GNNs, is validated using intra-validation, inter-validation, clinical application, and extra-validation. The datasets and models used are clearly shown in Table 2. Matplotlib 3.5.3 was used to plot the ROC and PR curves, and Seaborn 0.12.0 was used to create violin plots.

**Table 2.**
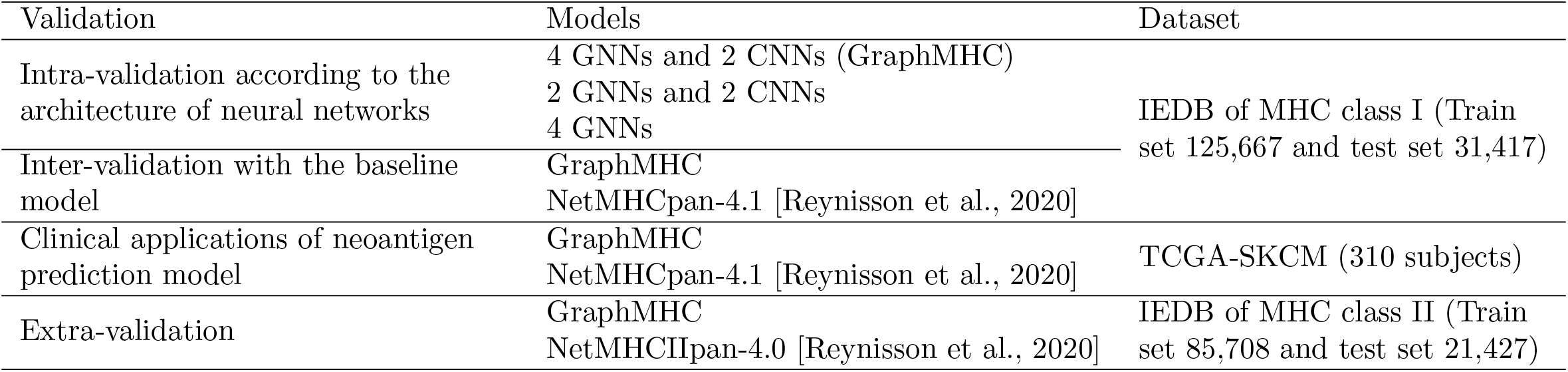
Four methods used for GraphMHC model validation: intra-validation, inter-validation, clinical application, and extra-validation.

### Intra-validation: Combining graph convolution with convolution improves prediction accuracy

The comparison results obtained for the different models according to the layer configuration of the neural network are shown in the center columns of Table 3. The model comprising 4-layer GNNs and 2-layer CNNs, GraphMHC, shows the highest performance. Analysis of the models comprising 2-layer GNNs and CNNs and only 4-layer GNNs indicated that feature extraction is effectively performed in the CNNs layers.

**Table 3.** Intra-validation according to model architecture of neoantigen classification and inter-validation using the baseline model.

| Metric | 4 GNNs and 2 CNNs (GraphMHC) | 2 GNNs and 2 CNNs | 4 GNNs | NetMHCpan-4.1 |
| --- | --- | --- | --- | --- |
| AUC-ROC | **0.922** (0.919-0.925) | 0.872 (0.869-0.876) | 0.688 (0.683-0.693) | 0.904 (0.900-0.907) |
| Sensitivity | **0.884** (0.881-0.888) | 0.839 (0.834-0.843) | 0.586 (0.580-0.591) | 0.834 (0.830-0.838) |
| Specificity | 0.810 (0.805-0.814) | 0.746 (0.742-0.751) | 0.690 (0.685-0.695) | **0.943** (0.941-0.946) |

Abbreviations: GNNs, graph neural networks; CNNs, convolutional neural networks; AUC-ROC, area under receiver operating characteristic curve. Values in parentheses indicate 95% confidence interval. Boldface typefaces represent the highest values among methods.

### Inter-validation: Graph-based model shows better sensitivity than string-based model

The results of comparing GraphMHC with the baseline model, NetMHCpan-4.1, in the rightmost column of the Table 3 indicate that GraphMHC perform well in terms of AUC-ROC and sensitivity and badly for specificity and *F*_1_-score. These results indicate a high rate for true positives and a low rate for false negatives, suggesting the production of meaningful results for fatal diseases such as cancer.

### Clinical applications: Groups divided by the graph-based model are clinically discriminated

The most common method of using the median as the threshold value for dividing groups was inherited in this study. [McGranahan et al., 2016, Kim et al., 2018, Ghorani et al., 2018] Examination of the 5-year survival on the upper right in Figure 3A indicates that the differences between groups divided using GraphMHC are borderline (*p*=0.061). On the other hand, the results obtained using NetMHCpan-4.1 were not significant in any of the observation periods, aligning with the results of previous studies in which no significance has been reported (*p*=0.567) for 10-year survival under this method. [Ghorani et al., 2018] Comparison of the biomarker scores in two groups in Figure 3B show that significant differences were obtained for stromal scores when using GraphMHC, but not for other cases.

**Fig 3.**
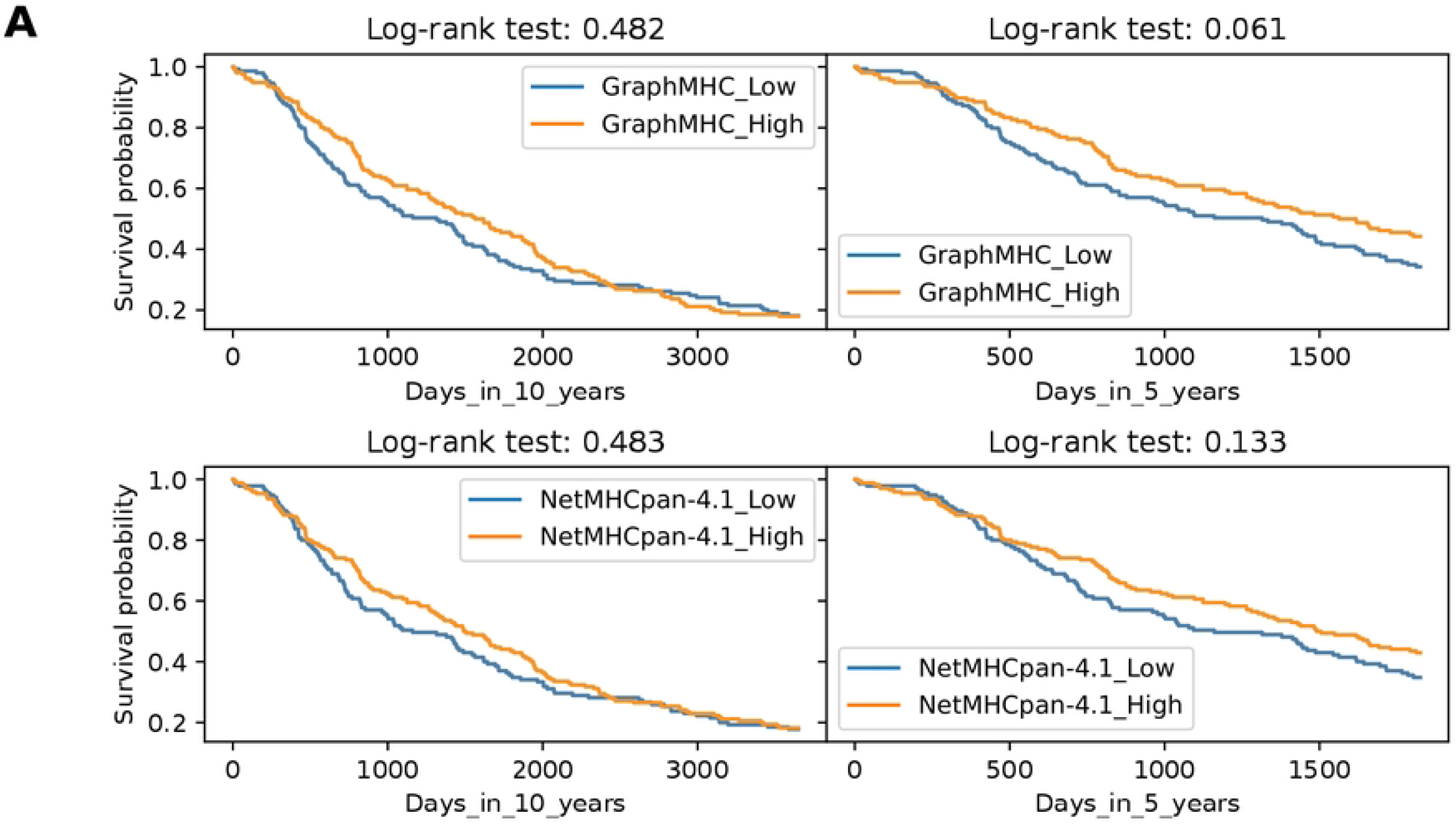
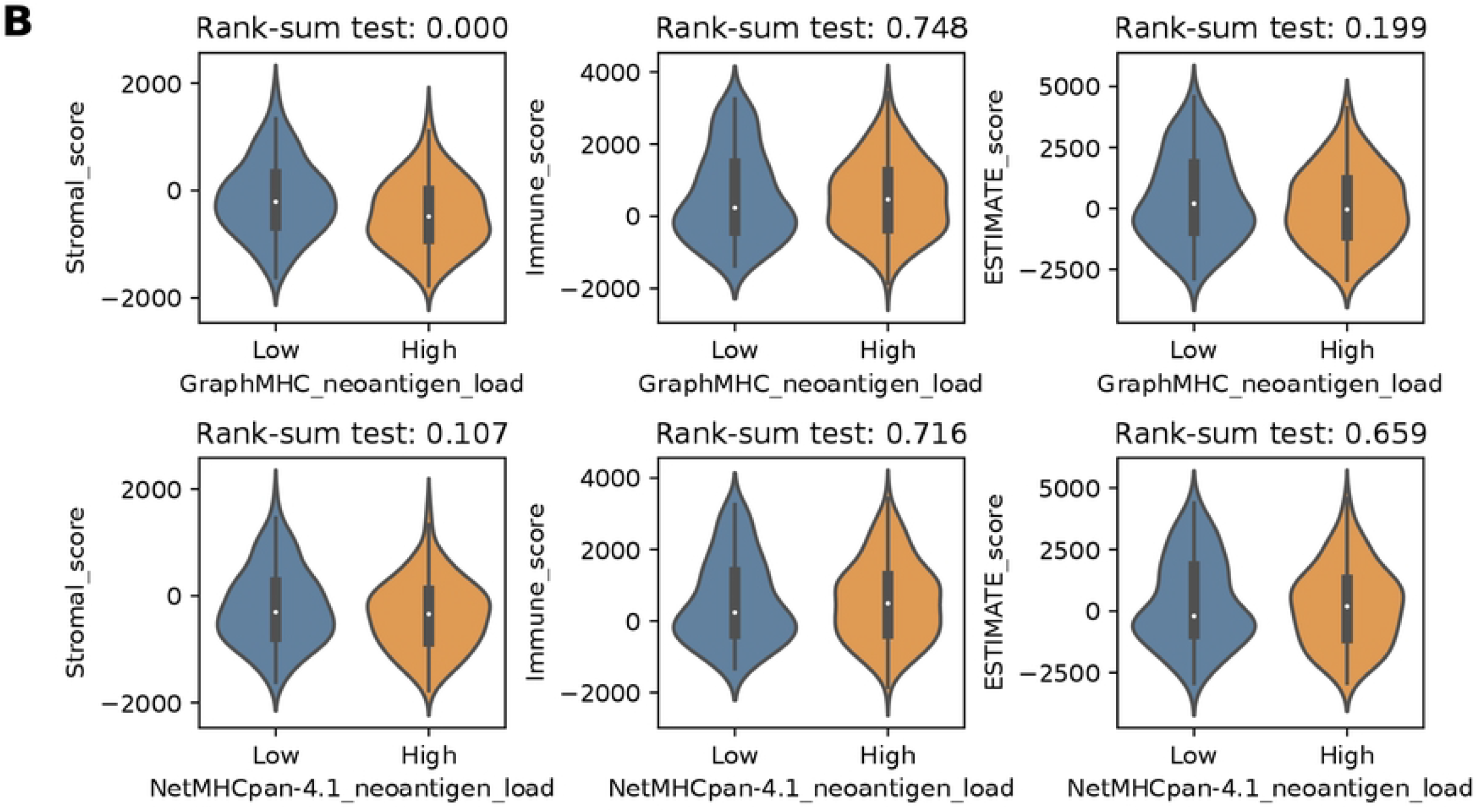
Comparison of high and low neoantigen load groups using TCGA-SKCM data. **A** Comparison of overall survival in high and low groups according to the median number of loaded neoantigens. The upper row is the result of grouping by GraphMHC while the lower row was obtained using NetMHCpan-4.1. Examination of the 5-year survival on the upper right indicates that the differences between groups divided using GraphMHC are borderline. **B** Comparison of biomarker scores in high and low groups according to the median number of loaded neoantigens. The upper row is the result of grouping by GraphMHC while the lower row was obtained using NetMHCpan-4.1. Comparison of the stromal scores in two groups show that significant differences were obtained when using GraphMHC.

### Extra-validation: GraphMHC model shows the best performance for most metrics in MHC class II data

The following are the evaluation results of the test set of 21 427 samples obtained from the MHC class II datasets in the IEDB. Comparison of GraphMHC and the corresponding baseline model NetMHCIIpan-4.0, applied to MHC class II data in Table 4, confirms GraphMHC showed higher performance in terms of AUC-ROC and sensitivity, as indicated by the results of inter-validation. Other scores showed similar or slightly lower values. The *F*_1_-score was the same as the value derived using the baseline model, and the specificity obtained was low.

**Table 4.** Extra-validation applied to MHC class II data.

| Metric | GraphMHC | NetMHCIIpan-4.0 |
| --- | --- | --- |
| AUC-ROC | **0.874** (0.869-0.878) | 0.834 (0.829-0.839) |
| Sensitivity | **0.813** (0.808-0.818) | 0.788 (0.782-0.793) |
| Specificity | 0.770 (0.764-0.776) | **0.798** (0.792-0.803) |

Abbreviations: AUC-ROC, area under receiver operating characteristic curve. Values in parentheses indicate 95% confidence interval. Boldface typefaces represent the highest values obtained using different methods.

## Discussion

### Implications of the neoantigen prediction study using the GNN

This study is the first to predict binding by modeling the biochemical molecular structure using information that describes MHC protein sequences and peptide sequences as candidate neoantigen. It is noteworthy that feature extraction and binding modeling are possible only when the inherent structural information from the sequence data is used and no additional external information is included. In other words, no separate interaction mechanism or through calculation is required; the structure and connection information between nodes and edges in the graph structure itself are aggregated and extracted through layered automatic feature extraction, and the minute attractive and repulsive forces that occur between complex and diverse atoms are simulated when using this method. As a similar research case, a protein folding model using GNN with reduced parameters from AlphaFold 2 [Jumper et al., 2021] has been reported to simulate the interaction between MHC and peptides, [Delaunay et al., 2022] indicating the possibility of using GNN for the structural prediction of MHC-peptide binding.

In addition, the differences in the research results obtained in previous studies are not limited to the performance of the model. Application of the proposed model, GraphMHC, to clinical data suggested that it has better discrimination compared to those observed previously. These results suggest that the GraphMHC model can be used as a biomarker for both cancer metabolism and the cancer immune response.

### Potential application suggested from research results

This research model can be used in biological experiments or for medical prediction or prevention. One case in which binding affinity was evaluated using NetMHCpan in *in vitro* immunoprecipitation and liquid chromatography-tandem mass spectrometry experiments(LC-MS/MS) [Khan et al., 2022] and one in which the CD8^+^ epitope was predicted and validated using NetMHC *in vivo* can be cited as close examples. [Duarte et al., 2015] In terms of screening, it is possible that accurate information about suitability can be obtained via sequencing for patients who are to receive immune checkpoint inhibitors, enabling patient-specific treatment. The model can also be used as a biomarker, dividing groups for response to cancer immunotherapy, and for comparison between two groups. [Kim et al., 2020, Abbott et al., 2021] Even more noteworthy is its possible use in preventing cancer by using it in the development of peptide vaccines that can activate T cells. [Sahin et al., 2017, Ott et al., 2017, Zahm et al., 2017] On the other hand, the results of this study could also be used to predict similar amino acid bindings such as the SARS-CoV-2 (COVID-19) antigen. [Prachar et al., 2020, Li et al., 2021] Furthermore, the application of this model could be considered not only for amino acids, but also for modeling ligand or drug binding that can be represented by SMILES. From these various perspectives, this study can be considered a pioneering breakthrough in precision medicine research.

### Limitations of the study

Despite the original suggestions and potential applications, this study also has some limitations. The information in the IEDB concerning the binding affinity between MHCs and peptides, including HLA-type polymorphisms is incomplete, although the experimental data are steadily accumulating. Inevitable uncertainties are also associated with the conversion and transformation of data via several different methods. In this regard, attempts to call variants more accurately using deep learning have been reported. [Poplin et al., 2018] In addition, there is no guarantee that the conversion of the 34-mer amino acid sequence corresponding to the polymorphic residues in the HLA types referenced to by NetMHCpan is absolute. In terms of the model itself, the internal structure of the GNN model means that it occupies more memory than the string-based neural network model, indicating that a server-side service would be useful. Beyond the neoantigen load, several other points require consideration when expanding the research scope of immunotherapy. Even in cases where large numbers of neoantigens are loaded, the prognosis is often poor. Therefore, not only the foreignness of a tumor as an antigenic mutation, but also the tumor sensitivity to mutations that are exported from the inside to the outside of a cancer cell should be considered and different machine learning methods applied. [Blank et al., 2016, Kim et al., 2020] It is also necessary to consider TCR binding, [Fritsch et al., 2014] for which other machine learning methods are being developed. [Tung and Ho, 2007, Tung et al., 2011, Lu et al., 2021]

### Research topics not included in this study

Since this study aimed to extract intrinsic information from a given amino acid sequence, the research methodology of using the extracted dataset with additional data was not included. This is because the methodologies used in studies reporting in this field are not as yet verifiable because of the small number or the artificial nature of datasets used. For example, one model that used molecular information about amino acids as well as binding affinity, Neopepsee, [Kim et al., 2018] reached high prediction accuracy with an AUC of 0.981 by applying a support vector machine (SVM) model that combined the binding affinities from the IEDB with parameters such as immunogenicity, sequence similarity, and amino acid pairwise contact potentials. [Saethang et al., 2013] The study was limited by the small sample size, with only 311 positive epitopes and 14 633 mutant negative peptides included, and the IMMA2 dataset used to calculate the amino acid pairwise contact potentials in this study was composed of only 558 immunogenic and 527 non-immunogenic peptide values, [Tung et al., 2011] Another example, NetMHCpan-4.0, is a multilayer perceptron model that outputs binding affinities and eluted ligands from mass spectrometry, reached approximately 0.98. [Jurtz et al., 2017] This study has a total of 85 217 entries, but the results are limited because the negative entries were artificially generated. Other models derived from it, such as MHCflurry [O’Donnell et al., 2020] and DeepHLApan [Wu et al., 2019], have similar methods.

## Conclusion

The GraphMHC model predicts neoantigens by converting MHC protein and peptide binding to graphic structure using the intrinsic features of the sequences themselves.

The GraphMHC model, which is based on the GNN, showed high accuracy with a low false negative rate for predicting neoantigens as compared to the baseline model. A significant difference was also observed when using the GraphMHC model to divide data into two groups for testing clinical discrimination. The GraphMHC model can thus be considered suitable for predicting the response prognosis of immune checkpoint inhibition.

## Supporting information

**S1 Table. SMILES representation of MHC and peptides**.

**S1 Fig. Molecular structures of a MHC protein and a peptide, with upper part corresponding to the peptide and lower part corresponding to the MHC protein**.

**S2 Table. Measurement statistics for all MHC-peptide graphs**. Based on the binding affinity provided by the IEDB, non-binding is defined as IC_50_ ≤ 500 nM and binding as IC_50_ *>* 500 nM. Statistics are expressed from the median (the 1st quartile - the 3rd quartile).

**S2 Fig. Graph structure of a MHC protein and peptide**. MHC protein and peptide chains are composed using disconnected graphs.

**S3 Fig. Selection of subjects from TCGA-SKCM data**.

## Acknowledgments

The authors thank the anonymous reviewers for their valuable suggestions.

## Data availability

All data used are publicly available: The MHC class I dataset, http://tools.iedb.org/static/main/binding_data_2013.zip; the MHC class II dataset, http://tools.iedb.org/static/download/classII_binding_data_Nov_16_2009_prediction_scores.tar.gz; TCGA-SKCM data, http://firebrowse.org/; HLA types, https://static-content.springer.com/esm/art%3A10.1186%2Fs12920-020-0694-1/MediaObjects/12920_2020_694_MOESM3_ESM.xlsx; immunity scores, https://bioinformatics.mdanderson.org/estimate/tables/skin_cutaneous_melanoma_RNAseqV2.txt. The code of this research model is published on the following GitHub page: https://github.com/recognizability/GraphMHC

## Author notes

This research paper was published in February 2023 as H.J.’s doctoral dissertation.

